# Spalt-related is an integrated stress response-activated inhibitor of mTORC1-mediated growth

**DOI:** 10.1101/2025.09.19.677425

**Authors:** Onur Deniz, Ying Liu, Tuuli Kirkinen, Krista Kokki, Pau Clavell-Revelles, Jaakko Mattila, Ville Hietakangas

## Abstract

Anabolic and catabolic processes are coordinated by a conserved regulatory network, which includes the nutrient sensing protein kinase mTOR complex 1 (mTORC1) and the insulin- and stress-responsive transcription factor FoxO. In physiological setting these regulators align growth, storage, reproduction, and aging with nutrient availability. Here, we identify transcription factor Spalt-related (Salr), previously implicated in organogenesis, as a negative regulator of growth and lipid storage. In the *Drosophila* fat body Salr activates catabolic gene expression and restricts mTORC1-mediated cell growth. The genomic binding of Salr overlaps extensively with that of FoxO and similar convergence is observed between their mammalian homologs SALL1 and FOXO1. Both Salr and FoxO are activated upon fasting but respond to distinct cues: while FoxO displays transient activation and is responsive to AKT inhibition, Salr is activated in a slow and sustained manner through the integrated stress response. Once activated, Salr counters nuclear localization of FoxO. Together, Salr and FoxO are converging transcriptional activators of catabolism during nutrient stress.

## INTRODUCTION

Metabolic pathway activities are tremendously adaptive to adjust cellular and systemic homeostasis in response to external conditions, including nutrient availability. Specific tissues, such as liver in vertebrates and fat body in insects, serve as metabolic hubs which coordinate systemic homeostasis. This sets high requirements for signal integration, i.e. an ability to simultaneously read and combine information from multiple inputs to generate a coordinated metabolic output. Impaired integration of nutrient-sensing signals can lead to systemic pathophysiologies, such as metabolic syndrome and type II diabetes (Klein et al. 2022; Priest and Tontonoz 2019). Protein kinase mTOR complex 1 (mTORC1) is active when nutrients and energy are readily available and it promotes the activity of anabolic pathways (Goul, Peruzzo, and Zoncu 2023). mTORC1 activity is counteracted by the Forkhead box class O (FoxO) family of transcription factors, which activate catabolic genes (X. Xu et al. 2012) and inhibit mTOR activity by promoting the expression its negative upstream regulator Sestrin (J. H. Lee et al. 2010). FoxO regulation integrates information from multiple sources: it is inhibited by insulin-like signaling through AKT mediated phosphorylation (Brunet et al. 1999; Puig et al. 2003) and activated by several stress conditions, such as oxidative stress, heat and genotoxic insults (Chen et al. 2010; Hopkins, Ragsdale, and Seo 2025; Huang and Tindall 2007; Rodriguez-Colman, Dansen, and Burgering 2024; Teleman et al. 2008). Despite these insights, we still lack in full understanding of how signal integration in metabolic hub organs is achieved in physiological setting.

Spalt transcription factors are Zn-finger transcription factors with conserved regulatory roles in patterning and organogenesis (De Celis and Barrio 2009). Mutations in two members of the human Sal-like (SALL) transcription factor family cause syndromes with impaired organogenesis, namely Townes Brocks Syndrome (SALL1) and Okihiro Syndrome (Al-Baradie et al. 2002; Kohlhase et al. 1998). Their roles were originally discovered in *Drosophila*, where the two paralogs Spalt-major (Salm) and Spalt-related (Salr) regulate cell proliferation and survival downstream of Decapentaplegic (Dpp) signaling in the developing wing (De Celis, Barrio, and Kafatos 1996; Sturtevant et al. 1997). Beyond development and organogenesis, however, the physiological roles of Spalt-like transcription factors are poorly understood. Here we report an unexpected role of *Drosophila* Salr as a fasting activated inhibitor of growth and lipid storage in fat body cells. Salr is deeply integrated into the core nutrient sensing regulatory network, displaying striking and conserved similarities with FoxO in target gene binding profiles. Salr targets include components of the mTORC1, such as *Sestrin* and *PRAS40*, and Salr is sufficient and necessary to restrict the growth promoting activity of mTORC1 (Parmigiani et al. 2014; Wang et al. 2007). Salr and FoxO possess many similar functional properties, but they respond to distinct upstream cues. While FoxO is regulated by insulin-like signaling and displays rapid and transient activation upon fasting, Salr is activated through the integrated stress response (ISR) pathway during prolonged nutrient deprivation. Thus, our results imply that two converging pathways control catabolic gene expression upon fasting depending on its duration and the upstream signaling cues involved.

## RESULTS

### Inhibition of growth and lipid storage by Salr

Through systematic analysis of transcription factors, we discovered a novel growth inhibitory role for Spalt-related (Salr) in the *Drosophila* larval fat body. When overexpressed in the fat body (CG-Gal4) Salr was sufficient to delay larval growth (Figure 1A) and pupariation (Figure 1B). In addition to systemic growth regulation, Salr overexpression in fat body clones (Tubulin [flipout] Gal4) strongly reduced cell size demonstrating cell autonomous function (Figure 1C). The growth inhibition was accompanied by smaller nuclear size, reflecting inhibition of endoreplication, as well as reduced relative size of the nucleolus (Figure 1C). To overcome the early lethality of Salr full body mutants (Barrio et al. 1999) and to analyze the tissue autonomous role of Salr, we generated Salr CRISPR/Cas12a transgenic flies, simultaneously targeting multiple regions of Salr (Figure S1A). Consistent with the role as a growth inhibitor, nuclei in tissue-specific knockout fat body cells were significantly larger than in control animals (Figure 1D) and they displayed significantly increased relative nucleolar size (Figure 1D). These data were further confirmed by using fat body specific knockdown by Salr RNAi (Figure S1B, C). The observed changes in nucleolar size suggested that Salr regulates ribosome biogenesis and consequently protein biosynthetic capacity. This was indeed the case as evidenced by O-propargyl-puromycin (OPP) labelling, reflecting reduced and elevated levels of newly synthesized proteins in Salr overexpressing and knockout cells, respectively (Figure 1E, F).

**Figure 1.**
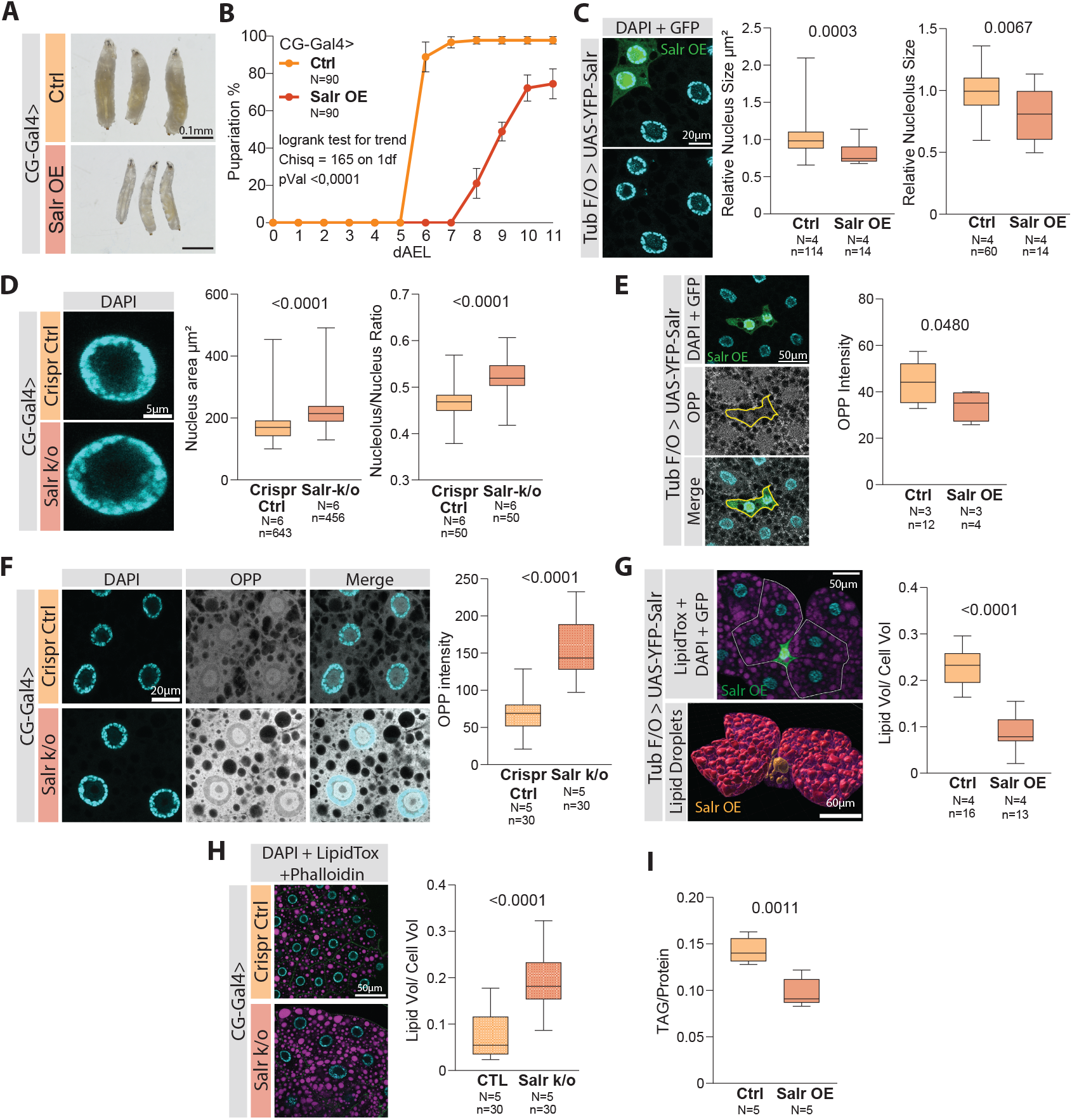
Salr inhibits cell growth and lipid storage cell autonomously. **A**) Representative image of early third instar larvae overexpressing Salr in the fat body. **B**) Fat body specific Salr overexpression leads to a delayed pupariation, dAEL: days after egg laying. **C**) Salr inhibits growth cell autonomously. Representative image of Salr overexpressing clones (Tubulin flip-out) marked with GFP and YFP. Quantification of nuclear and relative nucleolar sizes. **D**) Fat body specific CRISPR/Cas12a knockout of Salr leads to increased nuclear and nucleolar sizes. Representative image of Salr knockout and control nuclei stained with DAPI. Quantification of nuclear and nucleolar sizes. **E**) Salr overexpression decreases protein synthesis. Representative image of GFP and YFP marked Salr overexpressing clones stained with DAPI and OPP. Quantification of OPP intensity in Salr overexpressing and control cells. **F**) Salr CRISPR/Cas12a knockout in the fat body increases protein synthesis. Representative image of Salr knockout cells stained with DAPI and OPP. Quantification of OPP intensity of Salr knockout and control cells. **G**) Salr overexpression decreases lipid droplet volume in fat body cells. Representative image of GFP and YFP marked Salr overexpressing clone (Tubulin flip-out) stained with DAPI and LipidTox (above). 3D model of lipid droplets inside the white line (below). Lipid droplets of Salr overexpressing cell highlighted (orange). Quantification of total lipid volume to cell volume ratio in Salr overexpressing cells. **H**) Salr CRISPR/Cas12a knockout increases lipid droplet volume in fat body cells. Representative image of Salr knockout cells stained with DAPI, LipidTox and Phalloidin. Quantification of lipid droplet volume to cell volume ratio. **I**) Quantification of whole larval TAG levels, normalized to protein levels. P value and chi-square in **B** was calculated by logrank test for trend. P values in **C-I** were calculated with unpaired t-test with Welch correction. N = biological replicates, n = technical replicates.

As Salr overexpression in the fat body leads to smaller and thinner animals (Figure 1A), we explored the effects of Salr on lipid storage by staining fat bodies using the neutral lipid stain LipidTox. 3D analysis of confocal images revealed that Salr overexpressing clones have strongly decreased total lipid droplet volume compared to neighbouring control cells (Figure 1G). Conversely, fat body specific Salr knock-out increased total lipid droplet volume (Figure 1H). Moreover, fat body specific Salr overexpression reduced whole larval TAG levels (Figure 1I). In conclusion, Salr is necessary and sufficient to inhibit cell growth and lipid storage in the larval fat body.

### Salr controls nutrient responsive catabolic genes

As growth and lipid storage are closely controlled by nutrient availability, we wanted to test the possible nutrient regulation of Salr (Texada, Koyama, and Rewitz 2020). As a reporter, we used ectopic Salr with a C-terminal GFP inserted into the genomic locus (Kudron et al. 2018). Indeed, Salr displayed significantly elevated expression in the fat body of nutrient deprived larvae and was returned to control levels after refeeding (Figure 2A). The elevated Salr expression upon starvation was confirmed by qPCR analysis of endogenous Salr mRNA levels (Figure 2B). To explore the physiological role of Salr, we analyzed larval survival during starvation. Fat body specific Salr knockout displayed modestly but significantly impaired resistance towards starvation (Figure 2C). Similar starvation sensitivity was observed upon RNAi mediated knockdown of Salr in the fat body (Figure S2A). To obtain a global view of Salr downstream processes, we analyzed gene expression changes in the Salr gain- and loss-of-function fat bodies of early 3^rd^ instar non-wandering larvae. Comparison with published datasets on starvation-regulated genes revealed that Salr function mimicked the gene expression response observed in starved animals (Figure 2D, E) (Hasygar et al. 2021). Overrepresentation analysis revealed that Salr activates catabolic responses, including genes involved in autophagy and lipid catabolism (Figure 2F-H, Figure S2B-F). Conversely, Salr displayed a prominent inhibition of genes involved in mitochondrial ribosome biogenesis (Figure S2B, C, S2G-I).

**Figure 2.**
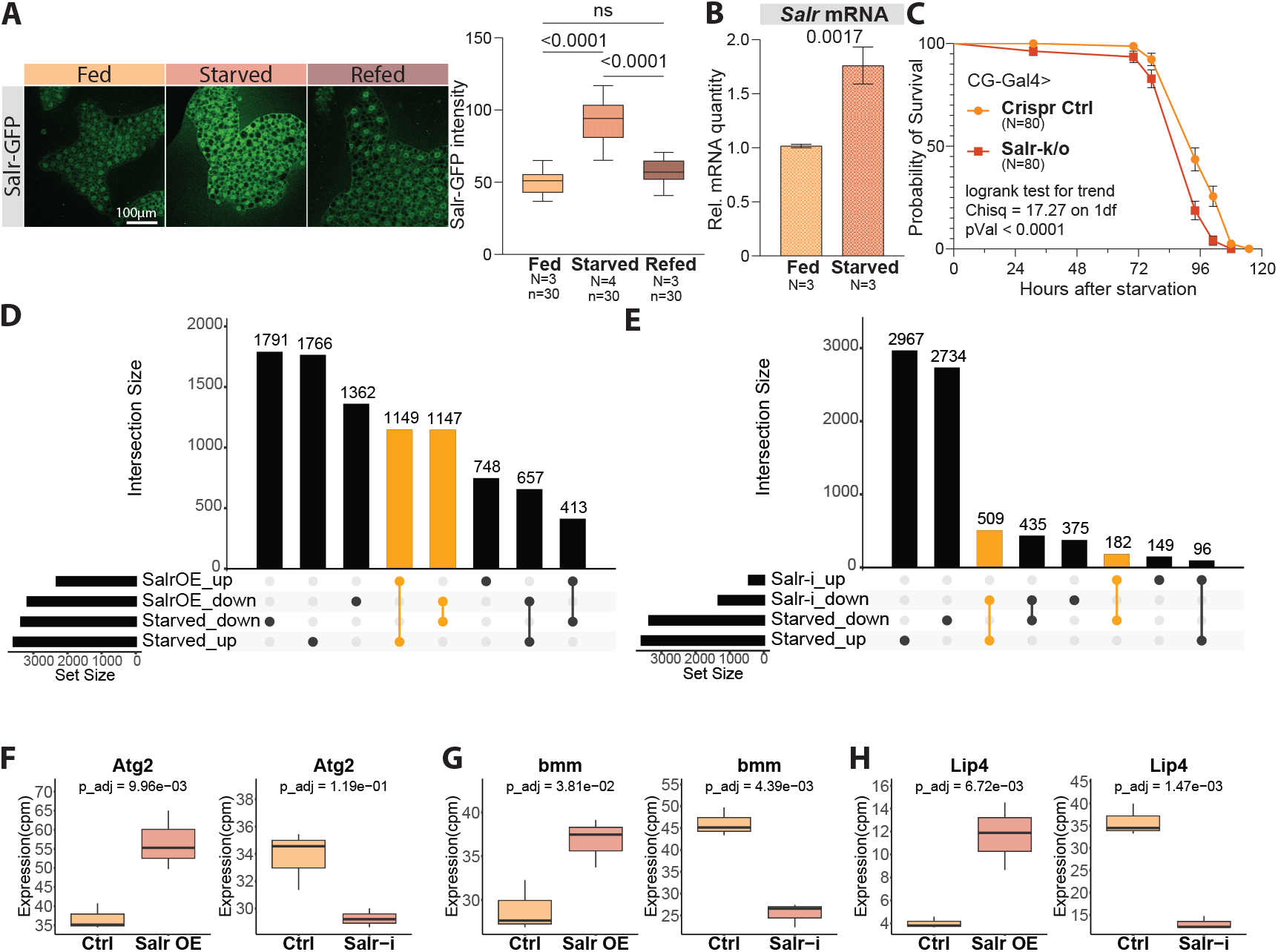
Salr is activated by fasting and activates catabolic gene expression. **A**) Endogenously GFP tagged Salr levels increase upon fasting and return to control levels after refeeding. Representative image and quantification of fat body cells with GFP tagged Salr under fed, 5h fasting and 6h refeeding. **B**) qPCR quantification of salr mRNA under fed and fasted conditions. **C**) Salr CRISPR/Cas12a knockout in the fat body leads to impaired survival under nutrient deprivation. **D & E**) UpSet plots of fasting induced differentially expressed genes (DEGs) and Salr gain- and loss-of-function DEGs. Highlighted intersections show fasting-mimicking gene regulation upon Salr gain- and loss-of-function. **F**-**H**) Salr activates catabolic genes, including those involved in autophagy and lipid catabolism. Atg2, bmm, and Lip4 expression from RNA-Seq datasets upon fat body specific Salr overexpression and knockdown. P values in **A** were obtained by one-way ANOVA followed by Tukey’s test with multiple comparison correction. P value in **B** was calculated by unpaired t-test with Welch correction. P value and chi-square in **C** was calculated by logrank test for trend. Adjusted P values in **F**-**H** were obtained in differential expression analysis of RNA-Seq with Benjamini–Hochberg correction. N = biological replicates, n = technical replicates.

### Salr inhibits mTORC1 signaling

To identify direct targets of Salr, we used the ModENCODE chromatin immunoprecipitation sequencing (ChIP-Seq) data from *Drosophila* white pupae (The modENCODE Consortium et al. 2010). Many of the genes in the Salr regulated pathways, including autophagy, lipid catabolism and mitochondrial ribosome biogenesis, were indeed direct targets (Figure S2J-L). Interestingly, among the Salr direct target genes were several prominent negative regulators of the mTOR complex 1 (mTORC1). These include *sestrin, pras40* and *reptor*, which were upregulated upon Salr overexpression and downregulated by Salr knockdown (Figure 3 A-D, Figure S3A-D).

**Figure 3.**
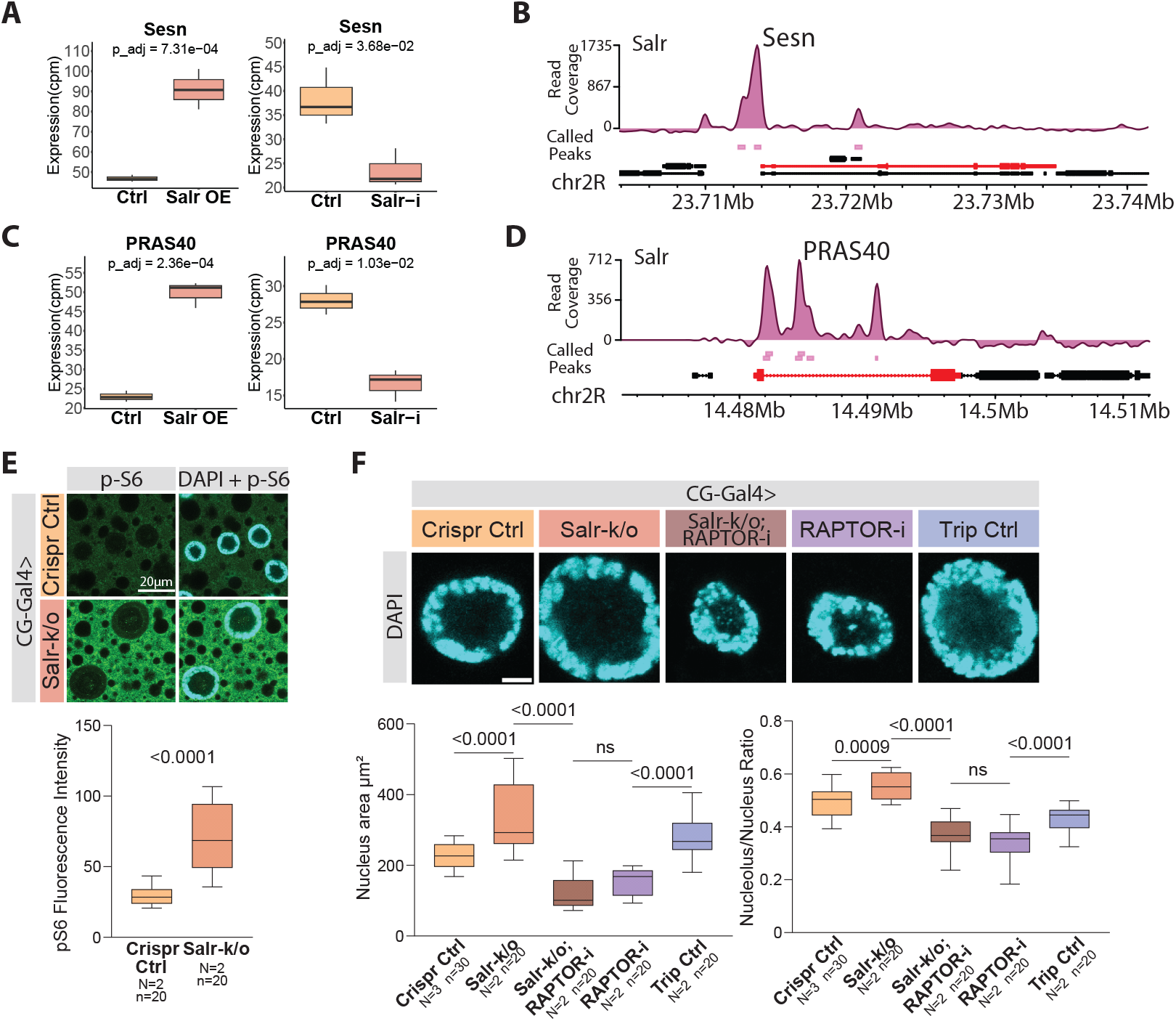
Salr inhibits growth by restricting mTORC1 activity. **A**-**D**) Salr targets and upregulates negative upstream regulators of mTORC1, Sesn and PRAS40. **A & C**) Sesn and PRAS40 expression from RNA-Seq datasets showing up- and downregulation upon fat body specific Salr overexpression and knockdown, respectively. **B & D**) Called peaks and track coverage of Salr ChIP-Seq on Sesn and PRAS40 loci (highlighted with red). **E**) Fat body specific Salr CRISPR/Cas12a knockout leads to elevated mTORC1 activity. Representative images and quantification of Salr knockout cells stained with DAPI and phospho-S6. **F**) Salr negatively regulates growth through mTORC1 signalling. Representative images of fat body nuclei of Salr CRISPR/Cas12a knockout in combination with Raptor RNAi. Quantification of nuclear and relative nucleolar sizes. Adjusted P values in **A** and **C** were obtained by differential expression analysis of RNA-Seq with Benjamini–Hochberg correction. P value in **E** was calculated by unpaired t-test with Welch correction. P values in **F** were obtained by one-way ANOVA followed by Tukey’s test with multiple comparison correction. N = biological replicates, n = technical replicates.

The gene expression data suggest that Salr promotes a gene expression program to suppress mTORC1 signaling. To test this hypothesis, we used fat body specific Salr CRISPR and monitored mTORC1 activity by antibody staining against phosphorylated ribosomal protein S6 (pS6), a well-established reporter of mTORC1 pathway activity (Isotani et al. 1999). Indeed, inhibition of Salr led to prominent increase in S6 phosphorylation consistent with increased mTORC1 activity (Figure 3E). To test if Salr-mediated growth regulation is mTORC1-dependent, we performed a genetic epistasis experiment by simultaneous inhibition of Salr and mTORC1 (Raptor knockdown; (Kim et al. 2002)). Our results show that inhibiting mTORC1 fully suppressed the Salr LOF induced increase in nucleus and nucleolus size (Figure 3F), further supporting the conclusion that Salr inhibits growth by restricting mTORC1 activity in the *Drosophila* fat body.

### Salr and FoxO share common direct targets

The nutrient responsiveness and target genes of Salr prompted us to explore its relationship with known regulators of nutrient responsive transcription. FoxO transcription factor is a well-known downstream target of insulin-like signaling and regulator of catabolic processes (Puig and Mattila 2011). A genome-wide comparison of chromatin binding revealed a striking overlap between FoxO and Salr targets: 89% of Salr target genes were shared with FoxO (Figure 4A). Common targets include *atg2* and *atg17, brummer* (*bmm*), *PRAS40* and *Sestrin (Sesn)* encoding components of autophagy, lipid catabolism and mTORC1 signalling pathway, respectively. In addition to targeting same genes, the local chromatin binding profiles of Salr and FoxO to the *atg2, PRAS40, bmm, Sesn, atg17* gene regions were nearly identical, implying close functional cooperation (Figure 4B, Figure S4A-C). To analyze the possible conservation of Salr and FoxO target gene regulation, we analyzed ChIP-Seq data derived from human HepG2 hepatocellular carcinoma cells (The ENCODE Project Consortium 2012). A noticeable overlap between binding profiles of Salr homolog SALL1 and FoxO homolog FOXO1 was observed: 63% of SALL1 targets were shared with FOXO1, while 95% of FOXO1 targets were common with SALL1 (Figure 4C). Moreover, human orthologs of the *Drosophila* Salr targets *atg2* (ATG2A), *PRAS40* (AKT1S1), *atg17* (RB1CC1), *Sesn* (SESN3, SESN1) showed highly similar binding profiles for SALL1 and FOXO1 (Figure 4D, FigureS4D, E). This data suggests that the regulation of common target genes by Salr and FoxO is conserved in mammals.

**Figure 4.**
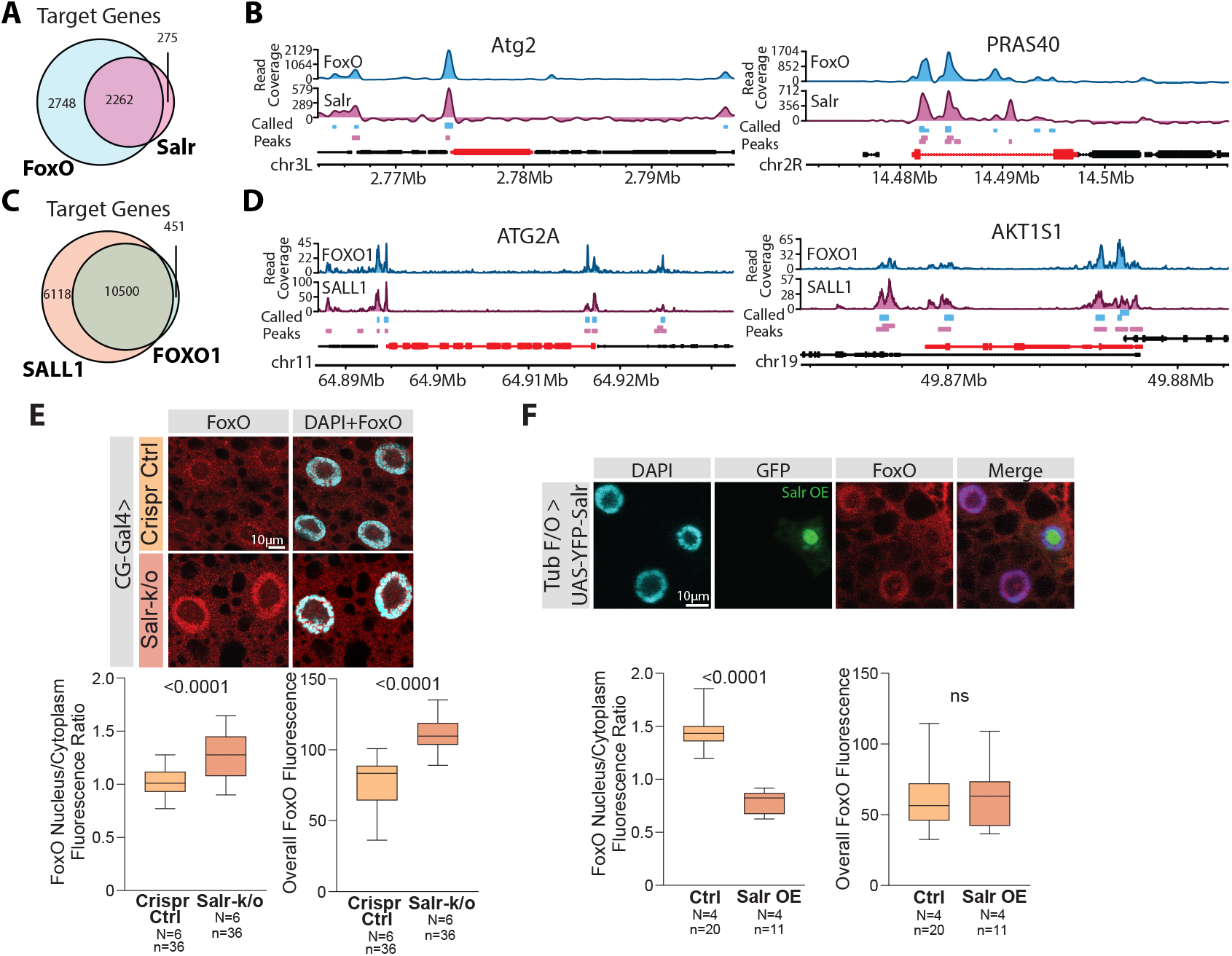
Salr shares common targets with and antagonizes FoxO. **A**-**D**) Salr and FoxO share common direct targets in Drosophila and human. **A**) Venn diagram of direct targets of FoxO and Salr. **B**) Called peaks and coverage tracks of Salr and FoxO ChIP-Seq on Atg2 and PRAS40 loci (highlighted in red). **C**) Venn diagram of direct targets of human SALL1 and FOXO1. **D**) Called peaks and coverage tracks of human SALL1 and FOXO1 on ATG2A and AKT1S1 loci (highlighted in red). **E & F**) Salr antagonizes nuclear localization of FoxO. **E**) Representative image of fat body specific Salr CRISPR/Cas12a knockout stained with anti-FoxO antibodies. Quantification of nuclear FoxO and overall FoxO levels. **F**) Representative image of Salr overexpressing clone stained with anti-FoxO antibodies. Quantification of nuclear FoxO and overall FoxO levels. P values in **E** and **F** were calculated with unpaired t-test with Welch correction. N = biological replicates, n = technical replicates

To investigate the interplay between Salr and FoxO functionally in *Drosophila*, we monitored intracellular localization of endogenous FoxO in Salr loss- and gain-of-function fat body cells. Surprisingly, FoxO nuclear localization correlated negatively with Salr expression. Upon fat body specific Salr loss-of-function FoxO showed increased nuclear localization and increased overall FoxO levels (Figure 4E). In contrast, Salr gain-of-function led to reduced nuclear/cytoplasmic ratio of FoxO without affecting the FoxO protein levels (Figure 4F). In sum, Salr binds to common genomic regions with FoxO and functionally antagonizes its nuclear localization.

### Salr is activated by prolonged fasting though the integrated stress response pathway

The antagonistic role of Salr on nuclear FoxO led us to postulate that while these factors target a common set of genes they might be activated by distinct physiological stimuli. Consistent with this, knockdown of Akt, a transducer of insulin-like signaling and a negative regulator of FoxO, had no effect on Salr expression (Figure 5A). Moreover, the activation dynamics of Salr and FoxO during starvation differed: while FoxO nuclear localization was increased by acute (2 h) fasting, Salr expression was highly activated only after prolonged fasting (16 h) (Figure 5B). Interestingly, the nuclear FoxO localization was significantly inhibited upon prolonged fasting, consistent with the observed inhibition of nuclear FoxO by Salr expression (Figure 5B).

**Figure 5.**
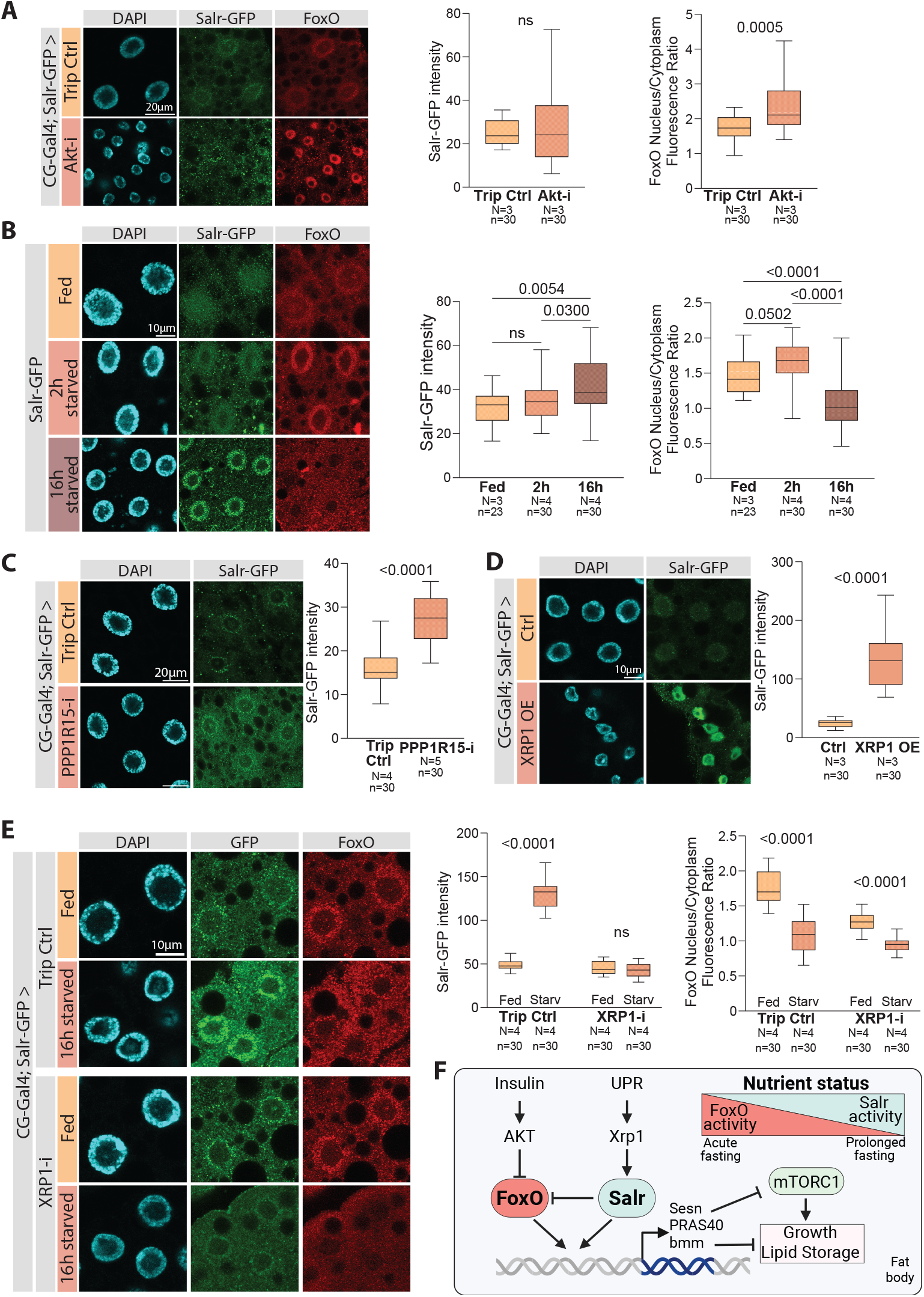
Prolonged fasting activates Salr through the integrated stress response pathway. **A)** Akt knockdown has no effect on Salr protein levels, while promoting nuclear localization of FoxO. Representative images of fat body specific Akt knockdown stained with anti-FoxO antibodies and anti-GFP antibodies for Salr-GFP. Quantification of Salr-GFP and nuclear versus cytoplasmic FoxO ratio. **B**) Differential activation dynamics of FoxO and Salr upon fasting. Representative images of fat body cells stained with anti-FoxO antibodies and anti-GFP antibodies for Salr-GFP after 2h and 16h fasting. Quantification of Salr-GFP and nuclear versus cytoplasmic FoxO ratio. **C**) Activation of the integrated stress response (ISR) pathway by PPP1R15 knockdown in the fat body activates Salr. Representative image of fat body specific knockdown of PPP1R15 with anti-GFP for Salr-GFP. Quantification of Salr-GFP intensity. **D**) Overexpression of XRP1 in the fat body increases Salr protein levels. Representative image of cells overexpressing XRP1, stained with anti-GFP for Salr-GFP. Quantification of Salr-GFP intensity. **E**) XRP1 knockdown prevents Salr activation upon prolonged fasting, but does not affect FoxO. Representative image of fat body specific XRP1 knockdown under fed and 16h fasted conditions, stained with anti-FoxO antibodies and anti-GFP antibodies for Salr-GFP. Quantification of Salr-GFP and nuclear versus cytoplasmic FoxO ratio. **F**) A model outlining Salr as a catabolic switch activated by prolonged fasting. P values in **A** and **D** were calculated with unpaired t-test with Welch correction. P values in **B** and **E** were obtained by one-way ANOVA followed by Tukey’s test with multiple comparison correction. N = biological replicates, n = technical replicates.

How is Salr activated upon prolonged starvation? Chronic nutrient shortage is known to activate the integrated stress response (ISR) leading to elevated phosphorylation of eIF2α (Ryoo 2024). The ISR is counteracted by phosphatase PPP1R15 (GADD34), which dephosphorylates eIF2α (Malzer et al. 2013). Knockdown of PPP1R15 led to significantly elevated Salr expression, supporting the idea of ISR-mediated regulation of Salr (Figure 5C). One of the downstream effectors of ISR is transcription factor Xrp1 (Brown et al. 2021), which we found to display chromatin binding close to the *salr* locus (Figure S5A). Overexpression of Xrp1 in the fat body led to a dramatic growth inhibition and larval lethality (not shown). Even transient overexpression of Xrp1 led to substantial fat body hypotrophy, which was accompanied by strongly elevated Salr expression (Figure 5D). We further tested if Xrp1 is necessary to activate Salr upon starvation, and, indeed, knockdown of Xrp1 fully abolished the elevated Salr expression by prolonged fasting (Figure 5E). In contrast, Xrp1 knockdown did not prevent the reduction of nuclear/cytoplasmic ratio of FoxO localization (Figure 5E). Collectively these data are consistent with a model that Salr and FoxO are activated by nutrient deprivation with differential kinetics and controlled by distinct upstream cues.

## DISCUSSION

Our study provides evidence for two functionally converging stress response pathways that activate catabolic gene expression and restrict the activity of anabolic mTORC1 signaling (Figure 5F). These pathways, controlled by Salr and FoxO transcription factors, respond to distinct upstream signals and are activated upon fasting with differential kinetics, FoxO displaying fast and transient activation and Salr being activated upon prolonged nutrient deprivation. Their functions appear mutually exclusive, as high expression of Salr is sufficient to counteract nuclear FoxO. ChiPseq data revealed high degree of overlap between the binding profiles, but future studies employing high-resolution binding site analysis are needed for more detailed insight for the molecular interplay between Salr and FoxO in the target promoters. The observed role for Salr as a regulator of metabolism and cell growth may appear surprising, regarding its well-establishes role in organogenesis (De Celis and Barrio 2009). However, similar examples exist, including Cabut, a regulator of Dpp signaling in wing patterning, which also controls metabolic target genes as a downstream effector of intracellular sugar sensing and the circadian clock (Bartok et al. 2015; Bejarano et al. 2008).

Our data shows that Salr functions downstream of ISR and Xrp1 and it restricts mTORC1 activity in fat body cells. Xrp1 is a known regulator of cell competition in developing *Drosophila* wing, being activated by proteotoxic stress and promoting the ‘loser’ status of cells destined to be eliminated (Baillon et al. 2018; C.-H. Lee et al. 2018; Recasens-Alvarez et al. 2021). Interestingly, inhibiting mTORC1 activity can alleviate the elimination of cells with proteotoxic stress through stimulation of autophagy and allowing clearance of misfolded proteins (Recasens-Alvarez et al. 2021). Considering the observed role of Salr as a downstream effector of Xrp1 as well as a negative regulator of mTORC1 and activator of autophagy genes, it will be interesting to learn whether Salr contributes to proteotoxicity-induced cell competition in the wing imaginal disc.

ChIPseq data derived from human hepatocellular carcinoma (HCC) cells revealed pervasive overlap in genome-wide binding profiles of SALL1 and FOXO1, including the binding to promoters of mTORC1 upstream regulators, which suggest conservation of the observed functions in humans. FOXO1 is a well-known tumor suppressor, and its expression is often reduced in HCC, which is associated with poor prognosis (Shi et al. 2018). Interestingly, SALL1 was also recently shown to function as a tumor suppressor in HCC (Saito et al. 2025). Considering that mTORC1 is a major driver in HCC, being deregulated in about 50% of cases (Yu et al. 2019), our findings raise the intriguing possibility that SALL1 and FOXO1 functionally converge to restrict mTORC1 activity in hepatocytes, thereby suppressing HCC tumorigenesis.

## MATERIALS AND METHODS

### *Drosophila* stocks and husbandry

Fly stocks used in this study: *w*^1118^ (BDSC 6326), UAS-YFP-Salr (Sánchez et al. 2010), Salr-GFP (BDSC 66443), UAS-Salr-RNAi (BDSC 29549), UAS-Dicer2 (BDSC 24650), UAS-XRP1 (FLYORF F003459 (Bischof et al. 2013)), UAS-XRP1-RNAi (BSDC 34521), UAS-Raptor-RNAi (BDSC 31529), UAS-FoxO-RNAi (BDSC 32427), UAS-Akt-RNAi (BDSC 33615), UAS-PPP1R15-RNAi (VDRC 107545), UAS-Luciferase-RNAi (BDSC 31603), UAS-EGFP (BDSC 6874), CG-Gal4 (BDSC 7011), UAS-Salr-crRNA (this study), UAS-Cas12a (this study), UAS-pCFD8 (this study).

Flies were maintained at 25°C, on medium containing agar 0.6% (w/v), malt 6.5% (w/v), semolina 3.2% (w/v), baker’s yeast 1.8% (w/v), nipagin 2.4%, propionic acid 0.7%.

To generate mosaic clones, larvae with the genotype HsFLP/+; UAS-YFP-Salr/UAS-GFP; Tub<Gal80, y^+^<Gal4/+ were heat shocked at 37×C for 12 mins at 24 h after egg laying.

### CRISPR/Cas12a-mediated tissue specific mutagenesis of Salr gene

Four guide RNAs targeting different locations of the Salr gene were designed ‘ggactcgtcagaagacctagatccgatctcgatcacgatccctgtaaatttctactaagtgtagatgtccggcggtcgggaatcactgcaatttctactaagtgtagatcttccgcaaacattacaaacgaaaatttctactaagtgtagatct taaccattggcttgctgagataacagggtcttcgttagctcg’ and cloned into the pCFD8 plasmid (#140619 AddGene) as described in (Port, Starostecka, and Boutros 2020). The Salr gRNA-pCFD8 plasmid were inserted into the ZH-51C (2R 51C1) site. An empty pCFD8 plasmid was inserted into the same location to be used as a control for CRISPR experiments. UAS-Cas12a plasmid (#140621 AddGene) was inserted into the attP40 (2L 25C7) site and the UAS-Cas12a and UAS-Salr-crRNA were recombined to be crossed with CG-Gal4 for fat body specific knockout of Salr.

### Nutrient deprivation of larvae

For fasting experiments, age matched L1 larvae were reared on standard laboratory food at equal density (30/vial) for 2 days. The larvae were then washed with PBS and transferred into 48 well plate containing starvation medium containing agar 0.5% (w/v) and erioglaucine dye 0.2% (v/v) for times indicated at figure panels. The plate was covered with film and small holes were opened for air circulation.

### Immunohistochemistry, LipidTox and OPP staining

For immunofluorescence staining, fat bodies were dissected from pre-wandering 3rd instar larvae (96h AEL) in PBS and fixed in 4% formaldehyde for 30 minutes. Tissues were washed with 0.3% Triton-X 100 in PBS and blocked in 2% NGS for 2 h. Subsequently tissues were stained with anti-Foxo (1:500) (Puig et al. 2003), anti-GFP (1:500) (Abcam), anti-phospho-S6 (1:500) (Cell Signaling) antibodies. LipidTox (1:400) (Thermo Fisher Scientific) staining was performed for fat bodies from early third instar larvae according to manufacturer’s instructions. Protein synthesis was evaluated using the Click-iT® Plus OPP Alexa Fluor 594 Protein Synthesis Assay Kit (Molecular Probes) as described in (Sanchez et al. 2016). Early third instar larvae were dissected in Shields and Sang M3 Insect Medium (Sigma) and fat bodies were stained in the Click-iT OPP Reagent at a 1:400 dilution (50 μM final concentration).

Samples were mounted in Vectashield mounting media with DAPI (Vector Laboratories) and imaged using the Leica SP8 microscope. Images were further processed by the ImageJ (NIH) and Imaris software.

### Image quantifications

For lipid droplet analysis, cell borders were defined across multiple z-sections using the Imaris custom surface tool. Lipid staining was confined to the rendered cell by masking the lipid staining within the rendered cell surfaces. The surface detection tool was then applied to the masked lipid staining to calculate total lipid volume within each cell, and this value was subsequently normalized to the corresponding cell volume. Nuclear size quantification was performed by using deep-learning based nuclear segmentation algorithm Stardist (Weigert et al. 2020). Nucleolar size quantifications were performed by manually selecting the region of interest (ROI) from images showing the maximum nucleus size. For the measurement of fluorescence intensities, background intensity was subtracted from mean intensity data from ROIs.

### RNA extraction, qPCR and mRNA sequencing

Age matched L1 larvae were collected and grown on standard laboratory food at equal density (30/vial). Fat body tissues of early third instar larvae were dissected, and RNA extraction was done using Nucleospin RNA II kit (Macherey-Nagel) according to the manufacturer’s instructions. For qPCR, cDNA was synthesized using a RevertAid H Minus First Strand cDNA Synthesis Kit (Thermo Scientific) following the manufacturer’s protocol. qPCR was performed with a Light cycler 480 Real-Time PCR System (Roche) using Maxima SYBR Green qPCR Master Mix (Thermo Scientific). The qPCR primers used were Salr ‘caagatctttggcagctactcggc’, ‘ccttcagattgcctttcgtggtg’ and Rp49 ‘agggtatcgacaacagagt’, ‘caccaggaacttcttgaatc’.

For mRNA-sequencing, libraries were generated using the TruSeq Stranded mRNA Library Prep Kit (Illumina) and subsequently sequenced with Illumina NextSeq500 technology to an average depth of 20 M reads per sample. The reads were mapped with TopHat to the *D. melanogaster* reference genome (FlyBase R6.10). The differential expression was quantified with limma package (v.3.28.8) (Ritchie et al. 2015) implemented in R/Bioconductor with Benjamini–Hochberg correction for adjusting p values. Genes with an adjusted p-value <0.05 were considered to be differentially expressed (DE). The genes with very low counts (cpm < 1 in more than one replicate per condition) were filtered. The raw sequencing data is deposited into the GEO repository (GSE123901). Genes identified as differentially expressed were subjected to over-representation analysis (ORA) to identify significantly enriched pathways. ORA was performed using the clusterProfiler package (v.4.12.6) (S. Xu et al. 2024) in R/Bioconductor. Pathways with an adjusted p-value < 0.05 with Benjamini–Hochberg correction were considered significantly enriched.

### ChIP-Seq

Browser Extensible Data (BED) files and control-normalized bigWig files of Salr (GSE257049), FoxO (GSE258971), SALL1 (GSE231113), FOXO1 (GSE170347) and XRP1 (GSE257114) were obtained from the ENCODE portal (https://www.encodeproject.org/). Peaks and coverage tracks were visualized with karyoploteR (v1.30.0) (Gel and Serra 2017). Promotor regions were defined as 500bp and 1kb upstream to the transcription start site for *D. melanogaster* and *H. sapiens* data, respectively.

## Statistical analysis

Statistical analyses were performed in GraphPad Prism 10 (GraphPad Software). For survival and pupariation rate analysis the Log-rank test was used. For parametric data, unpaired t-test with Welch correction or two-way ANOVA in conjunction with Tukey’s HSD test was used. The exact test, sample numbers and P values for each experiment is indicated in the figure legends. All quantitative data are shown as the mean ± SEM.

## Supporting information

All supplemental figures

## ACKNOWLEDGEMENTS

We thank Oona Kinnunen and Heini Lassila for technical assistance. Rosa Barrio, Norman Zielke, Bloomington *Drosophila* Stock Center, Vienna *Drosophila* Resource Center, Zurich ORFeome Project are acknowledged for fly stocks. This work was supported by Research Council of Finland (312439 to V.H.), Sigrid Jusélius Foundation (to V.H. and J.M.), Novo Nordisk Foundation: NNF19OC0057478 and NNF22OC0078419 (to V.H.), Jane & Aatos Erkko Foundation (to J.M.), Integrative Life Science Doctoral Program (to O.D.), Maud Kuistila Memorial Foundation (to O.D.), Ida Montinin Säätiö (to O.D.), Orion Research Foundation (to O.D.). The study was facilitated by the University of Helsinki *Drosophila* core facility (Hi-Fly), DNA Sequencing and Genomics Laboratory, and the Light microscopy unit (LMU) supported by Biocenter Finland and Helsinki Institute of Life Science.

## AUTHOR CONTRIBUTIONS

O.D.: investigation, project administration, formal analysis, visualization, conceptualization, writing - original draft, writing - review & editing, methodology, funding acquisition; J.L.: investigation, formal analysis, project administration, conceptualization; T.K.: investigation, formal analysis, visualization; K.K.: formal analysis; P.C.-R.: investigation; J.M.: supervision, writing - original draft, writing - review & editing, conceptualization, funding acquisition; V.H.: conceptualization, supervision, funding acquisition, writing - original draft, writing - review & editing, project administration.

## DECLARATION OF INTEREST

The authors declare no competing interests.

## Notes

### Competing Interest Statement

The authors have declared no competing interest.

