## Supplementary material for "Spalt-related is an integrated stress response-activated inhibitor of mTORC1-mediated growth": All supplemental figures

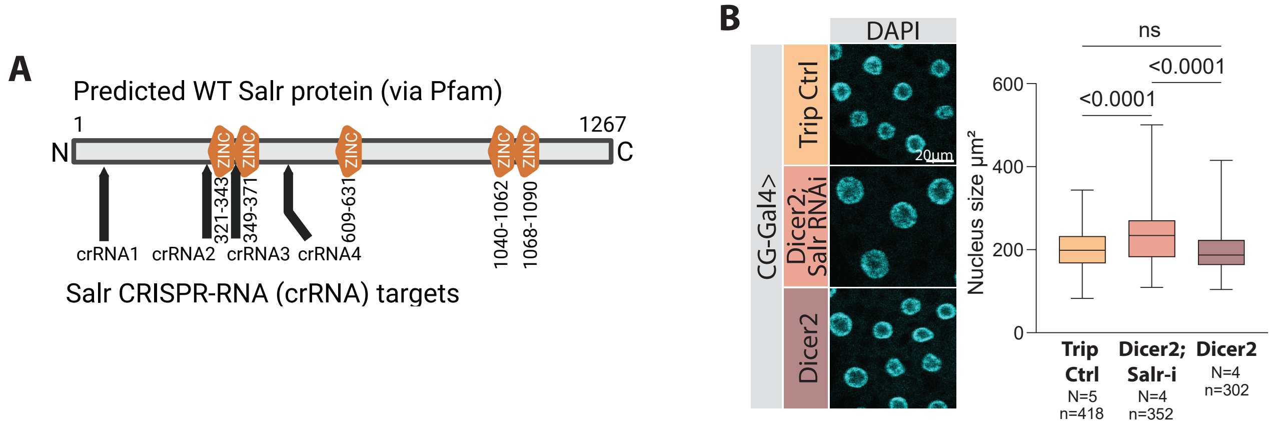


**Supplementary Figure 1.**

**A**) Salr CRISPR/Cas12a target sites on wild-type Salr protein. **B**) Salr knockdown with Dicer2 increases nucleus size. Representative image of fat body cells that are Salr knockdown with Dicer2, Trip control, and Dicer2, stained with DAPI. Quantification of nucleus size. P values in **B** were obtained by one-way ANOVA followed by Tukey's test with multiple comparison correction. N = biological replicates, n = technical replicates.

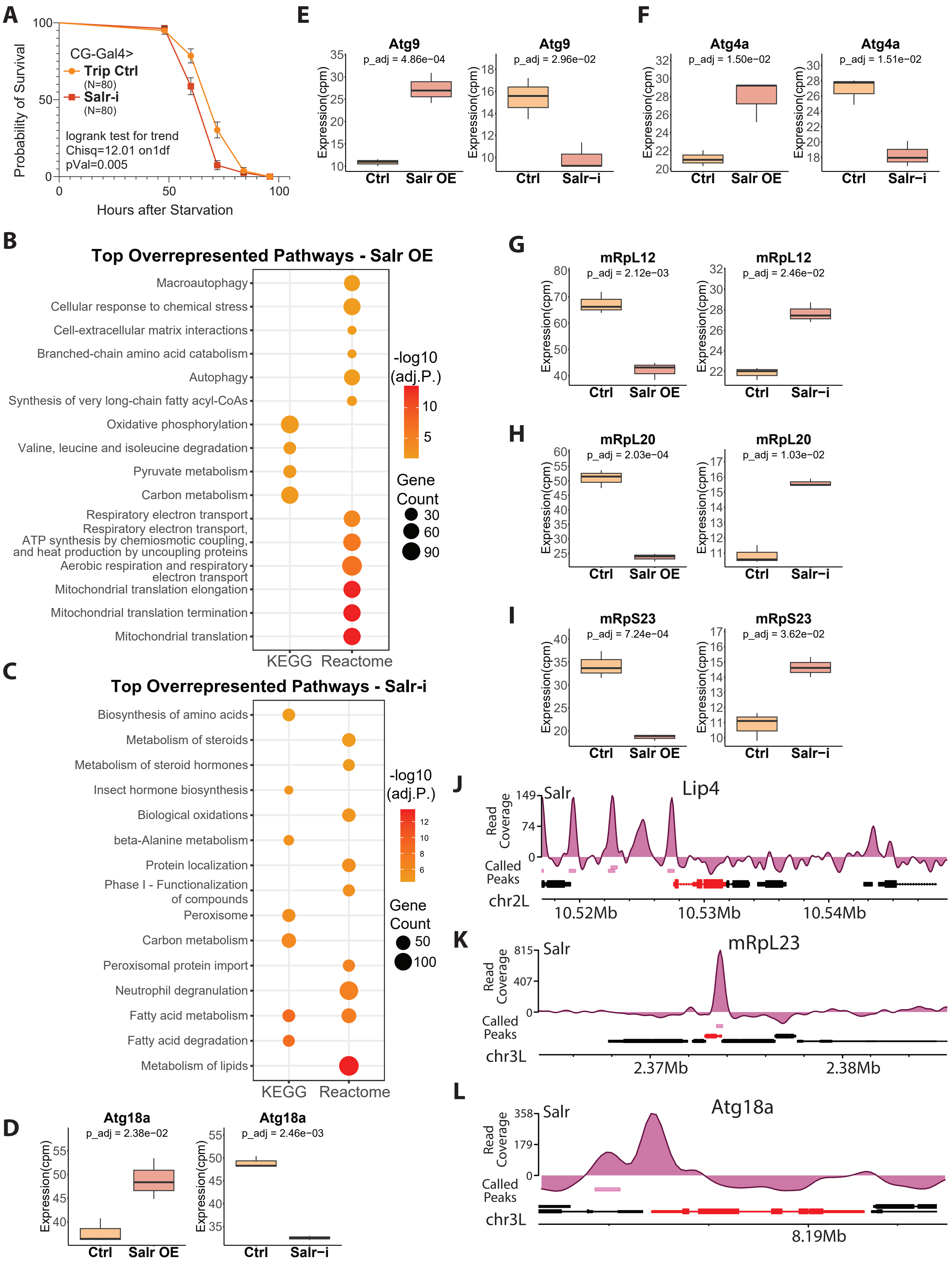


**Supplementary Figure 2.**
**A**) Salr knockdown in the fat body impairs survival under starvation. **B** & **C**) Unselected top overrepresented pathways from fat body specific Salr overexpression and knockdown RNA-Seq datasets, ranked based on their adjusted p value. **D-L**) Salr overexpression and knockdown in the fat body significantly regulates genes involved in lipid metabolism, autophagy, mitochondrial ribosome biogenesis, some of which are direct targets of Salr. **D-I**)

Atg18a, Atg9, Atg4a, mRpL12, mRpL20, mRpL23 expression from RNA-Seq datasets upon fat body specific Salr overexpression and knockdown. **J**-**L**) Called peaks and track coverage of Salr ChIP-Seq on Lip4, Atg18a, and mRpL23 genes highlighted in red. P value and chi-square in **A** was calculated by logrank test for trend. Adjusted P value in **B** and **C** were obtained with Benjamini–Hochberg correction. Adjusted P values in **D**-**I** were obtained in differential expression analysis of RNA-Seq with Benjamini–Hochberg correction. N = biological replicates.

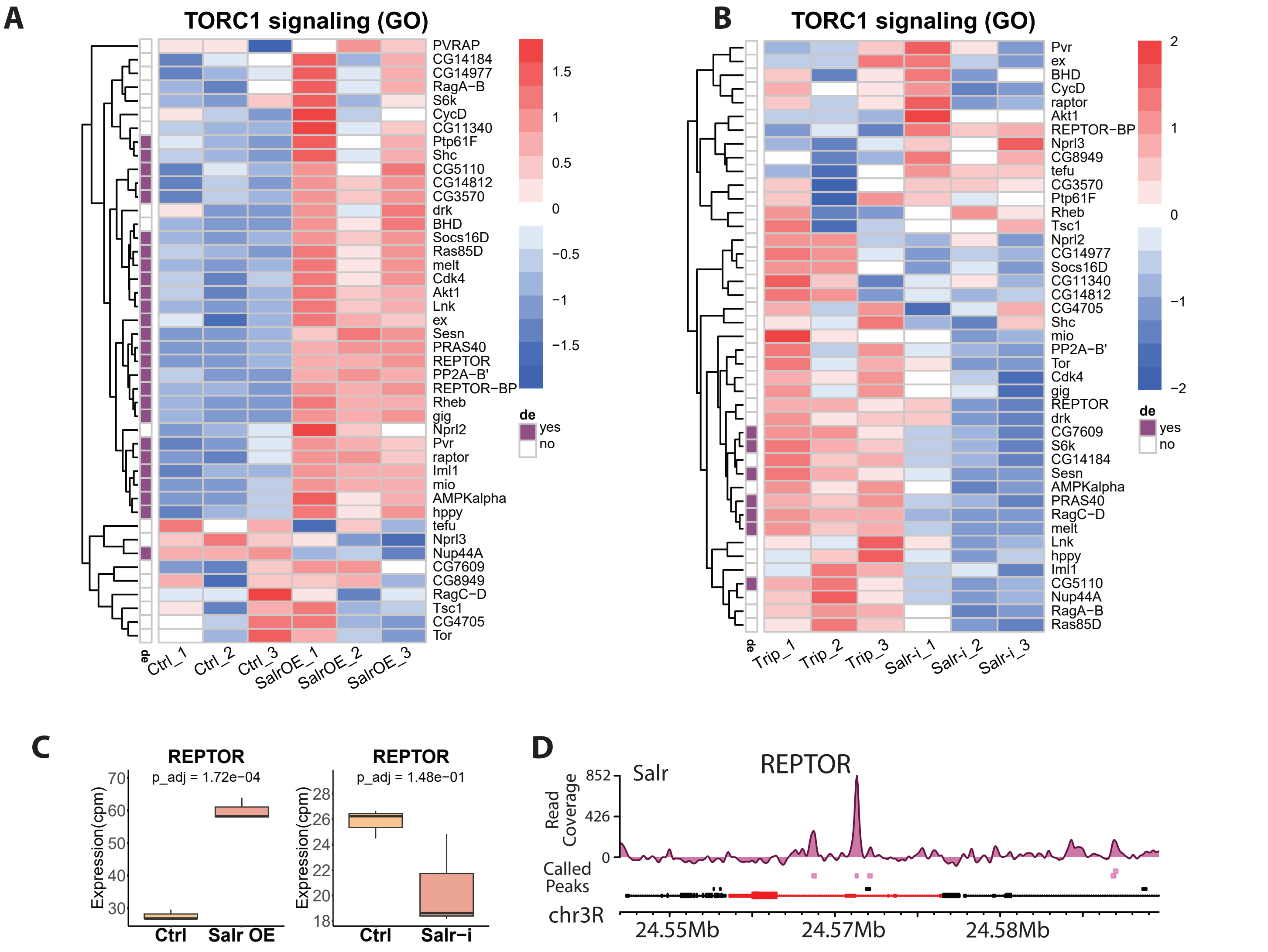


**Supplementary Figure 3.**
**A & B**) Salr regulates mTORC1 signalling pathway. Heatmaps of GO database defined mTORC1 signalling pathway from Salr overexpression and knockdown RNA-Seq datasets. **C & D**) Salr targets and regulates REPTOR expression. **E**) REPTOR expression from RNA-Seq showing up- and downregulation upon fat body specific Salr overexpression and knockdown, respectively. **F**) Called peaks and track coverage of Salr ChIP-Seq on REPTOR gene highlighted in red. Adjusted P value in **C** was obtained with Benjamini–Hochberg correction in differential expression analysis of RNA-Seq.

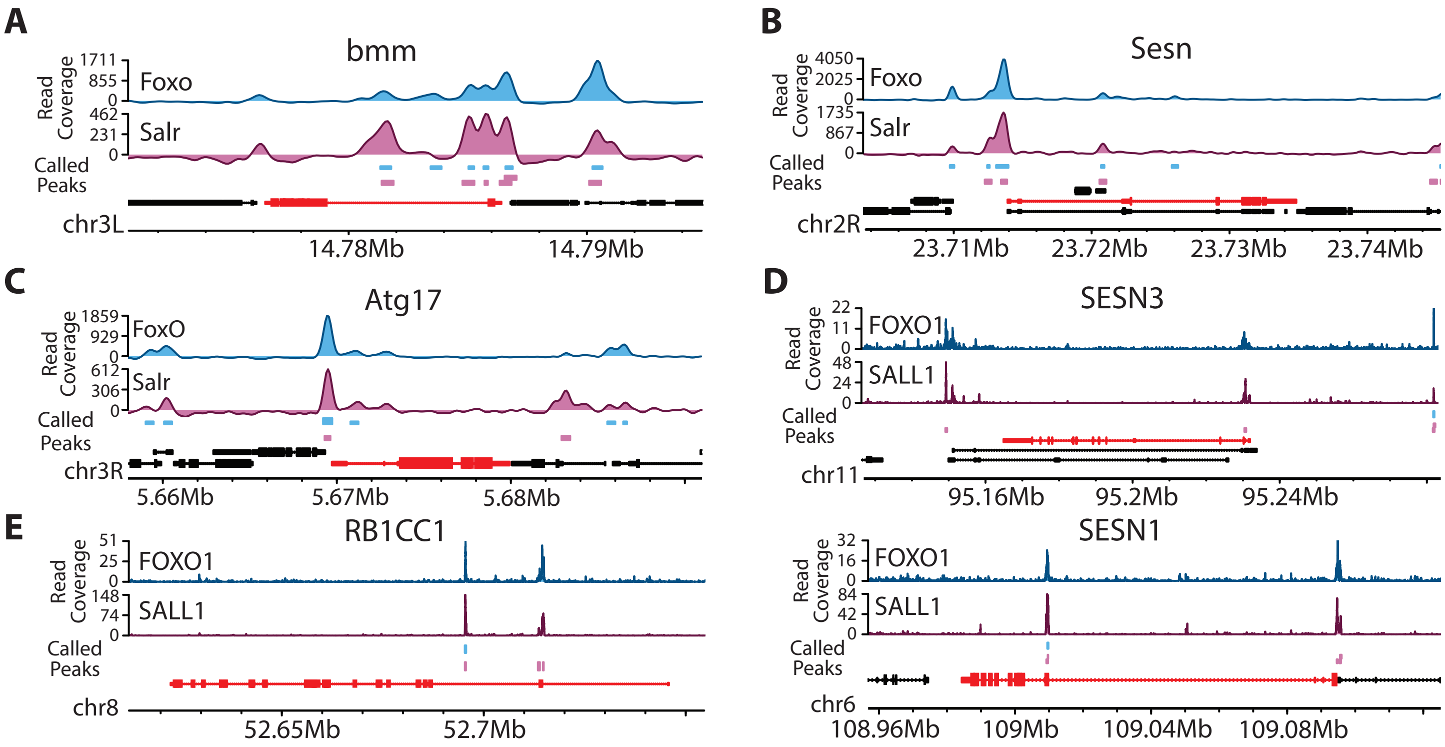


**Supplementary Figure 4.**
**A**-**E**) Common direct targets of Salr and FoxO in Drosophila are shared in humans. **A**) Called peaks and track coverage of Salr and FoxO ChIP-Seq on bmm. **B**) Called peaks and track coverage of Salr and FoxO ChIP-Seq on Sesn. **C**) Called peaks and track coverage of Salr and FoxO ChIP-Seq on Atg17. **D**) Called peaks and track coverage of SALL1 and FOXO1 ChIP-Seq on SESN3 and SESN1 (human ortholog of Sesn) **E**) Called peaks and track coverage of SALL1 and FOXO1 ChIP-Seq on RB1CC1 (human ortholog of Atg17).

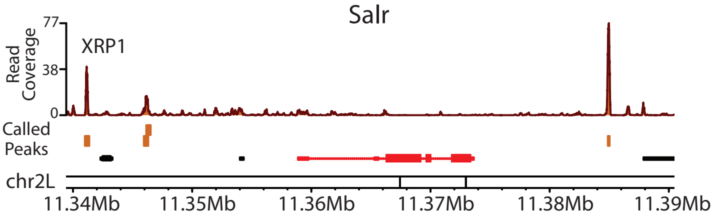


**Supplementary Figure 5.**
XRP1 targets Salr. Called peaks and track coverage of XRP1 ChIP-Seq on Salr gene which is highlighted with red.
